# The first chromosome-level annotated reference genome for *Chrosomus eos*, the northern redbelly dace

**DOI:** 10.1101/2023.12.21.572839

**Authors:** Ben Schultz, Elizabeth G. Mandeville

## Abstract

Hybridization amongst teleost fish species is an ongoing phenomenon with unclear implications. *Chrosomus eos*, the northern redbelly dace, is a species with an especially complex and fascinating history of hybridization with the closely related species *Chrosomous neogaeus*, the finescale dace. The two species historically hybridized during the last glacial maximum, forming a rare unisexual F1 hybrid lineage that utilizes gynogenetic reproduction. This hybridization event is currently thought to have only occurred during the glacial period, with subsequent dispersal of parents and hybrids into contemporary ranges during glacial recession and a cessation of further hybridization. All three maintain sympatric relationships, with limited evidence suggesting recent or contemporary hybridization between the parental species. To enable future research, a novel, high-quality, chromosome-level genome was produced through Dovetail Genomics and analyzed for completeness and the syntenic relationship between itself and the high-quality, well cited zebrafish *Danio rerio* genome.

## Introduction

The northern redbelly dace (*Chrosomus eos*, Cope 1862; formerly *Phoxinus eos*) is a small-bodied, freshwater Leuciscid minnow species, found in a wide and varied range across north-eastern North America. It has been extensively studied due to the unusual outcome of hybridization, or interbreeding with a related species, that occurred during glaciation. *C. eos*, along with the closely related finescale dace (*C. neogaeus*, Cope 1867), have formed an exceedingly rare phenomenon by producing an all-female, asexually reproducing hybrid. Across much of its range, *C. eos* is sympatric with *C. neogaeus*. Near the Last Glacial Maximum, estimated around 30,000–50,000 years before present, a hybridization event occurred between male *C. eos* and female *C. neogaeus* leading to the formation of an all-female asexual hybrid lineage that persists today alongside its parental species (Angers & Schlosser 2007, Binet & Angers 2005, Elder & Schlosser 1995). The contemporary range of *C. eos* and its hybrid is a result of dispersal into post-glacial habitats during the recession of the Laurentide ice sheet, extending as far west as the Canadian province of British Columbia, and spreading east along the 49th parallel through the Great Lakes region and St. Lawrence valley to Nova Scotia (Scott & Crossman 1973, Angers & Schlosser 2007). Asexual vertebrates are a rare phenomenon; less than 0.01% of described vertebrates are asexual, all of which arose by means of hybridization (Dawley 1989, Vrijenhoek *et al*. 1989). The asexual *Chrosomus* hybrids reproduce through gynogenesis, requiring the sperm of a male to stimulate embryological development while often not incorporating genetic material from the male (Goddard *et al*. 1998). Today, hybrid *C. eos-neogaeus* occur with both, or either one of the parent species in order to maintain their obligate gyongenetic relationship.

Previous work in this system suggested that there was no ongoing hybridization or back-crossing in *Chrosomus* dace, and that that all hybrid populations persisted as all-female asexual lineages from the initial hybridization event (Angers & Schlosser 2007, Mee & Taylor 2012, Schlosser *et al*. 1998). However, a recent study identified elevated diversity of hybrids, suggesting that there may be some evidence of contemporary hybridization (Monette *et al*. 2020), potentially connected with climate change, but this remains uncertain in part because of the limited resolution of previous genetic data. A large body of research has been performed on this hybridization complex using 10 DNA microsatellites, from which the majority of our understanding of the history of these fish is derived (Binet & Angers 2005, Angers & Schlosser 2007). By studying these species at increased genomic resolution, we aim to illuminate the genomic history of *C. eos, C. neogaeus*, and their hybrids, as well as gaining a better understanding of mechanisms of reproductive isolation and outcomes of hybridization. Using genomic data has improved our knowledge of other similar fish hybridization systems as in the gynogenetic Amazon molly, Poecilia formosa (Barbiano *et al*. 2013, Warren *et al*. 2018), where updated analyses suggested higher diversity than expected from gynogenesis alone, potentially as a result of backcrossing. Given the asexual reproductive strategy of *Chrosomus* hybrids, they offer insight into processes of speciation and the role of sex determining mechanisms in hybridization that their sexual parental species cannot, and it is therefore desirable to use high resolution genomic data to examine the evolutionary history of *Chrosomus* dace lineages in more detail.

Here, we present the first high-quality whole genome produced for *Chrosomus eos*, which we intend to use in further exploration of the history of hybridization between *C. eos* and *C. neogaeus*. We provide statistics and analysis of the overall structure and quality of the genome, and also explore the syntenic relationship between *C. eos* and the high-quality, well known zebrafish (*Danio rerio*) genome.

## Methods

### Fish collection and preparation

Field sampling occurred in October 2020 on private land in Melancthon, Ontario, Canada (UTM: 17N 563233 4882754; sampling conducted under University of Guelph Animal Use Protocol # 4443 and MNRF License to collect fish for scientific purposes # 10956456). Four *C. eos* were captured using a minnow trap and euthanized using a standard procedure of immersion into an MS-222 solution until overdose by anaesthesia achieves total opercular movement cessation. Standard length was recorded, and the fish were quickly dissected to parse out fin clips for DNA extraction and muscle tissue for high molecular weight DNA extraction and genome construction. These tissues, along with eye, upper digestive organs, and whole bodies were flash frozen in liquid nitrogen on-site and transported to a -80 freezer at the University of Guelph for holding.

### DNA extraction for identification

Fin clip tissue from the left pectoral fin was taken from each fish and extracted using a Qiagen DNeasy Blood and Tissue extraction kit, following manufacturer specified guidelines. Genetic identification using barcoding analysis, specifically the mitochondrial cytochrome oxidase subunit 1 (CO1) gene, confirmed each fish specimen was correctly identified as C. eos. The CO1 gene provides enough confidence to rule out the common hybrids due to it being inherited matrilineally - in a hybrid, we would expect the *C. neogaeus* mitochondrial genome (Angers & Schlosser 2007).

### Dovetail de novo assembly

Muscle tissue from two of the preserved fish were shipped to Dovetail Genomics LLC (Scotts Valley, CA) for de novo genome assembly. The Dovetail pipeline begins with high molecular weight DNA extraction. From this, continuous long-read libraries were prepared and sequenced using PacBio long read technology. Sequences were then assembled using `wtdbg2`, a fast and accurate long-read software tool (Ruan & Li 2020). To enhance the long-read assembly and generate a higher-quality genome, a second library was prepared using the proprietary Omni-C library preparation (a modified Chicago library preparation method Putnam *et al*. 2016) for short-read sequencing on an Illumina HiSeq X. These sequences were then fed into the HiRise software pipeline for scaffolding, creating highly accurate, long scaffolds. Concurrent to the production of the assembly, Dovetail performed a series of quality analyses including providing overall genome assembly statistics and BUSCO scores (see table 1). Once a high-quality final assembly was produced with a minimum single-copy BUSCO score of 70%, genome annotation was completed using RNA sequencing of flashfrozen tissues by Genewiz, with the Dovetail bioinformatics team completing the annotation. The annotation begins with repeat masking prior to ab initio gene prediction and annotation enhancement using RNAseq results (methods described in detail in Jarvis *et al*. 2017).

**Table 1:** Summary statistics describing the quality and completeness of the *C. eos* genome.

| Genome Assembly Statistics |  |
| --- | --- |
| Total Length | 1.2706 Gbp |
| Number of gaps | 1552 |
| Number of Scaffolds | 2344 |
| Longest Scaffold | 64894510 bp |
| N50 | 41636777 |
| L50 | 13 |
| BUSCO Score | 94.90% |
| Total Genome Masked | 54.98% |
| Class 1 TE repeats | 15.41 |
| Class 2 TE repeats | 20.42 |
| Total number of genes | 44326 |

### Syntenic Relationship with *Danio rerio*

We explored the syntenic relationship between *C. eos* and *D. rerio*, the most closely related model organism with a high quality genome, using Comparative Genomics’ (CoGe) synmap function (Lyons *et al*. 2008, Lyons & Freeling 2008). CoGe first uses Last, a variant of BLAST, to compare sequences from the genomes of each subject fish to a database (Frith *et al*. 2010). After then filtering out repeated matches, the sequences were fed into DAGchainer to identify syntenic genes (Haas *et al*. 2004). Finally, CoGe outputs a syntenic dotplot with colour coordinated points representing regions of synteny in both the same and inverted orientation. To further display the syntenic relationship between both genomes, we produced a circle plot using the program Circos (Krzywinski *et al*. 2009), which used the DAGchainer results from CoGe’s synmap output. To examine the phylogenetic relationships between the two species, we used the R package Fishtree (v 0.3.4) to produce a phylogenetic tree, and TimeTree 5 provided the estimates of divergence time (Chang *et al*. 2019, Rabosky *et al*. 2018a, Kumar *et al*. 2022). The zebrafish was chosen for use in this comparison because of the high-quality of its complete genome - it is the best reference genome of any reasonably close evolutionary relative - and common use of zebrafish as a model species.

## Results and Discussion

### Analysis of Final Assembly and Annotation

The final assembly contained 2344 scaffolds, with 25 chromosome-level scaffolds corresponding to the estimated haploid chromosome number of 25 (Dawley & Goddard 1988, Fig. 1). The total length of the genome was calculated to be 1270.66 Mb (Table), 23% less than the C-value estimated at 1.64, with the 25 chromosome level scaffolds accounting for 1101.85 Mb, 86.7% of the total size of the assembly (Dawley & Goddard 1988). The completeness of the final assembly was confirmed with a BUSCO score of 94.90%.

**Figure 1.**
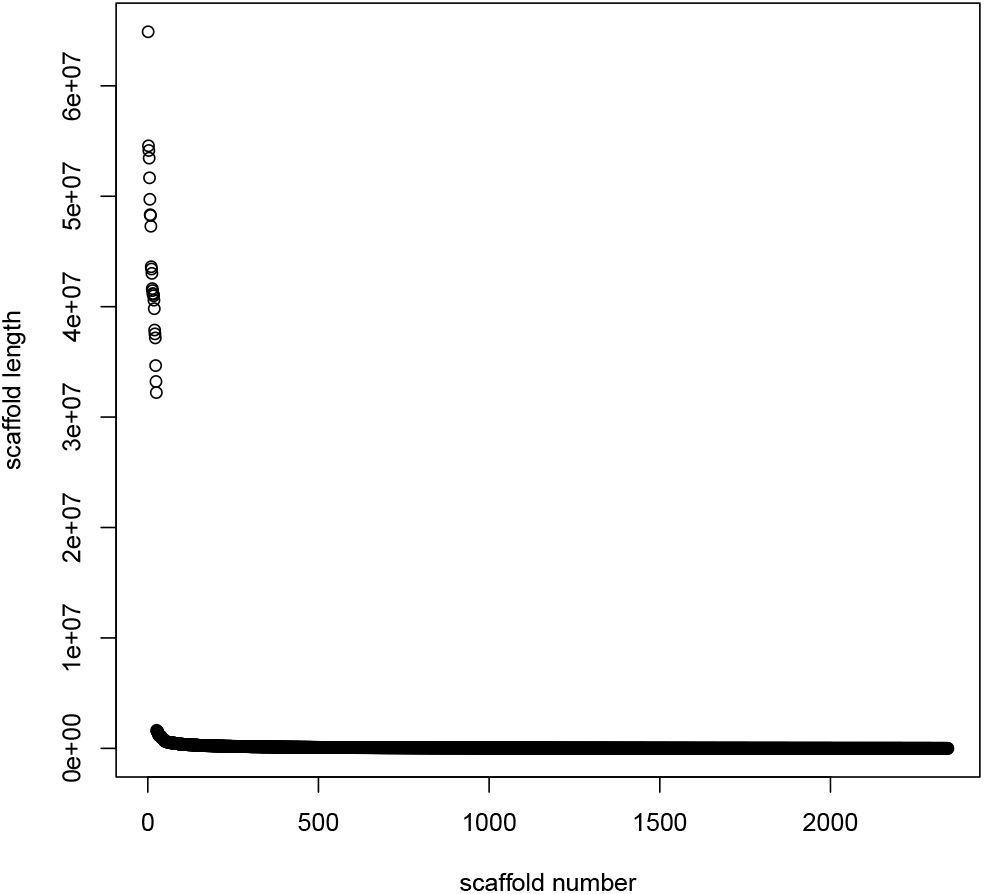
Plot showing the length of each scaffold by number; Critically, there is a substantial size difference between the 25th scaffold and the 26th, exemplifying the chromosome-level completeness of the new *C. eos* genome.

Repeat masking resulted in a total of 54.98% of the genome being masked, with Class 1 TEs repeats accounting for 15.41% and class 2 TEs repeats accounting for 20.42% (Table 1). Gene prediction resulted in a total number of 44,326 genes, while the coding region length was 66,456,841 bp.

### Synteny Between *C. eos* and *D. rerio*

The CoGe analysis provided DAGchainer results identifying over 25,000 regions of synteny between *C. eos* and *D. rerio*, spread across the genomes of both fish (Fig. 2). Overall, many regions of the *C. eos* genome maintain a syntenic relationship with at least one chromosome from *D. rerio*, with the exception of a large breakpoint between *C. eos* chromosome 5 and *D. rerio* chromosome 4 (Fig. 3 3). Insights provided by the syntenic relationship between *C. eos* and *D. rerio* allow us to key in on regions of interest within *C. eos*’ genome for further analysis, such as syntenic links between *D. rerio* chromosome 4, hypothesized to contain a zebrafish sex determining region (Nagabhushana & Mishra 2016), or areas of little synteny for identifying chromosomal rearrangements (Bhutkar *et al*. 2008). Given the high rate of transition between sex determining systems amongst fish, it is likely that this chromosome could vary greatly between both species (Bachtrog *et al*. 2014, Pennell *et al*. 2018). There is an estimated median divergence time between *C. eos* and *D. rerio* of 63 MYA, whereas the median time of divergence between *C. eos* and *C. neogaeus* stands at 17.9 MYA (Fig. 4; Rabosky *et al*. 2013, 2018b, Kumar *et al*. 2022).

**Figure 2.**
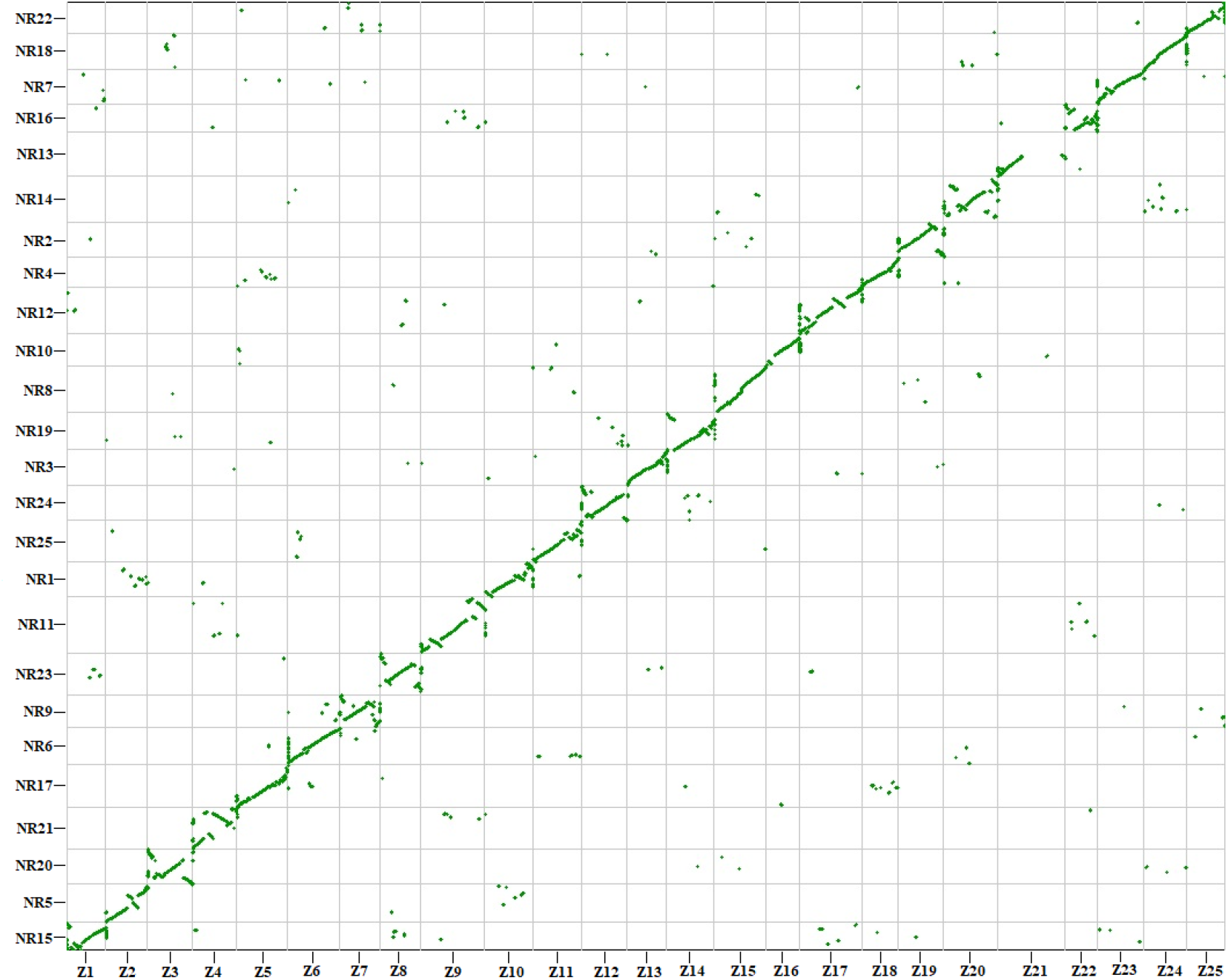
Comparative Genomics (CoGe) dotplot showing syntenic hits between the genomes of *C. eos* and *D. rerio*. The plot utilized the syntenic path assembly (SPA) function to arrange chromosome pairings in order of an unbroken syntenic relationship, using the *D. rerio* as the template on the x-axis. The *C. eos* genome is then assembled along the y-axis, with scaffolds positioned corresponding to which *D. rerio* chromosome they are syntenic with. In this orientation, break points and inversions are easily visually identified by their disruption of the linear relationship of the SPA.

**Figure 3.**
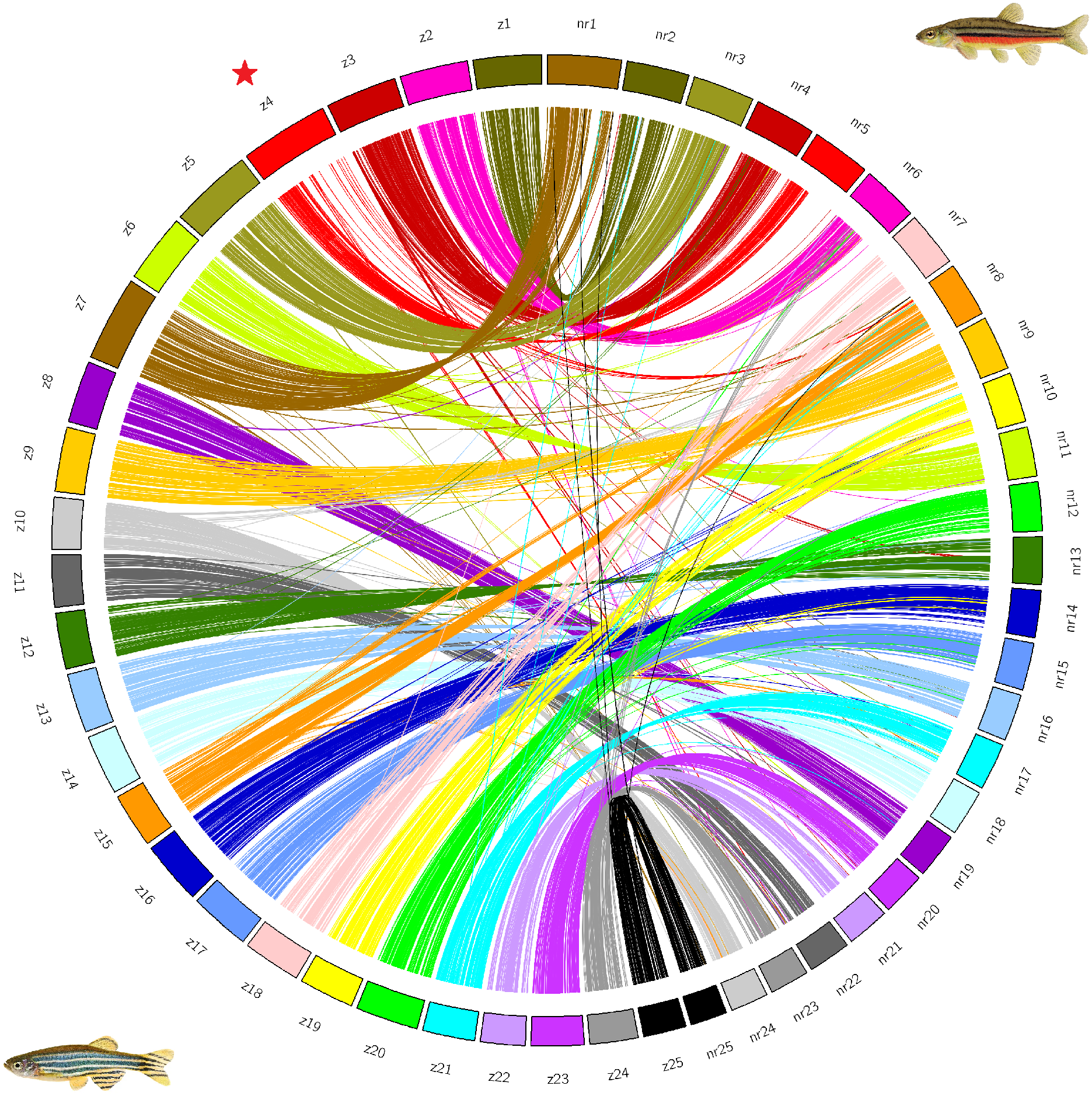
Circle plot produced using Circos showing syntenic links between the chromosomes of *D. rerio* on the left and *C. eos* on the right. The links and karyotype are coloured based on the syntenic relationship, using *D. rerio* as the template genome. To provide information on each syntenic link, the DAGChainer results from the CoGe dotplot were utilized.

**Figure 4.**
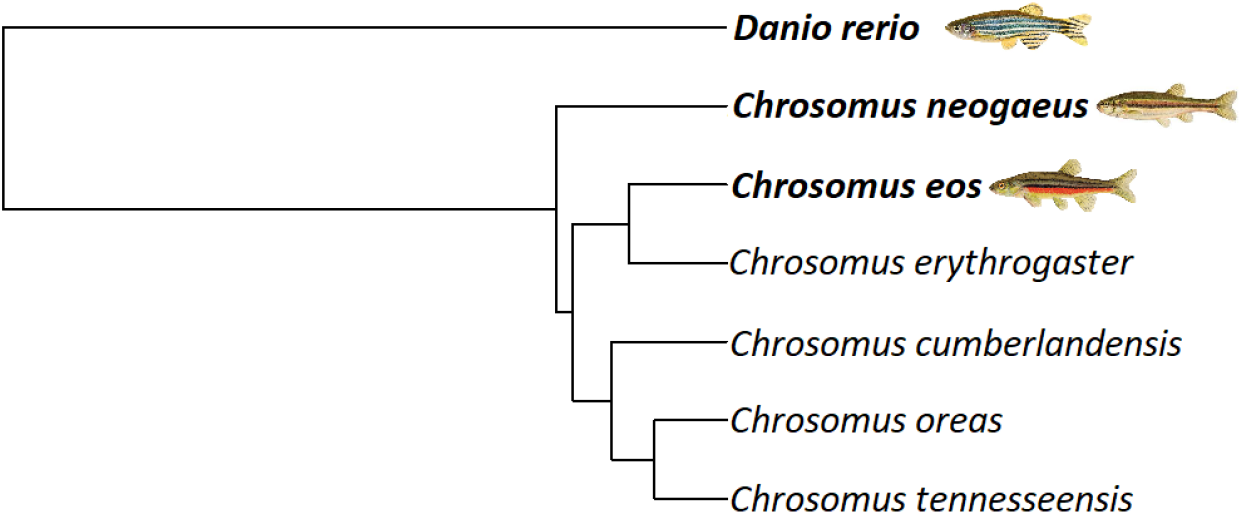
Phylogenetic tree including extant members of the *Chrosomus* genus, along with the distantly related *D. rerio. C. eos* and *C. neogaeus* diverged an estimated 17.9 million years ago, and *C. neogaeus* forms an outgroup from all other more closely related *Chrosomus* species.

Differential sex determination mechanisms between two species has frequently been a culprit for unequal hybridization outcomes (Orr 1997, Payseur *et al*. 2018), and in some known cases is associated with diversification within a clade (e.g. in sticklebacks; Natri *et al*. 2019). The method of sex determination in *C. eos* and *C. neogaeus* has been thus far undetermined, and it is unclear if the two *Chrosomus* species have shared or different mechanisms of sex determination. The presence of a unisexual hybrid suggests some potential genomic incongruence might prevent the formation of male first generation hybrids. Amongst Teleost fish sex determination varies greatly between taxa, including between closely related species of the same genus, with genetic, environmental, and hermaphroditic strategies occurring (Bachtrog *et al*. 2014, Sardell *et al*. 2020, Pennell *et al*. 2018). The utility of a complete reference genome provides the potential to discover previously cryptic sex determination mechanisms within *C. eos*.

This new high-quality reference genome will allow for the discovery of potential incompatibilities or genomic rearrangements between *C. eos* and *C. neogaeus* beyond the sex-determining regions of the genome as well. The presence of an asexually reproducing, all female hybrid lineage may suggest some level of genomic incompatibility between *C. eos* and *C. neogaeus*, potentially in keeping with Haldane’s rule, or potentially exposing Dobzhansky-Muller incompatibilities (Haldane 1922, Dobzhansky 1937, Muller 1942). Furthermore, the expansion of genomic data has the potential to increase the complexity of our understanding of the hybridization history between these two species, as has been seen in the Amazon molly (*Poecilia formosa*), where an F1 generation of hybrids was formerly thought to be frozen, until genomic evidence determined a history of backcrossing and recombination (Barbiano *et al*. 2013).

The production of a novel, high-quality reference genome for *C. eos* will enable a greater understanding of the evolutionary history and hybridization dynamics of not only *C. eos*, but all closely related taxa including *C. neogaeus*.

## Acknowledgements

We would like to thank J. Campbell, A.V. Meuser, A. Pitura, F. Tabak, and C. Pyne for assistance with fieldwork and bioinformatics. We thank D. McCauley and M. Kouprie for permission to sample fish in their stream. Genome sequencing was funded through an NSERC Discovery Grant and Launch Supplement to E.G. Mandeville. B. Schultz was also partially supported by a Research Support Grant from the Canada First Research Excellence Fund to E.G. Mandeville, through the Food From Thought program at the University of Guelph, and the Ontario Graduate Scholarship. Computing was accomplished using a Resources for Research Groups (RRG) computing allocation from the Digital Research Alliance of Canada.

## Author contributions

B. Schultz led field work, tissue sampling, data analyses, and writing. L. Mandeville contributed to study design, performed field work, coordinated initial genome sequencing, and contributed to writing and revising the manuscript.

## Data accessibility

Data and scripts used for analysis are publicly available on Github at: github.com/ bschultzuog/genome_code. This whole genome shotgun project has been deposited at DDBJ/ENA/GenBank under the accession JAZAVI000000000. The version described in this paper is version JAZAVI010000000. The raw reads from this project have been deposited with SRA and are available under the accession number PRJNA1054594.

